# miR-378a and NPNT coordinate autophagy regulation in podocytes through mTOR and MAPK signaling

**DOI:** 10.64898/2026.03.19.709781

**Authors:** Nina Sopel, Sarah-Maria Wangerin, Marie Hecker, Alexandra Ohs, Janina Müller-Deile

## Abstract

Autophagy is a critical homeostatic mechanism in podocytes, maintaining cellular integrity under stress and proteostatic challenges. Dysregulation of autophagy has been implicated in different glomerular diseases such as diabetes and membranous glomerulonephropathy (MGN), yet the underlying molecular drivers remain incompletely understood. We identified microRNA-378a (miR-378a), previously found upregulated in MGN, as a functional enhancer of autophagic flux in human podocytes and tubular epithelial cells. While miR-378a did not directly alter transcription of canonical autophagy genes (*ATG2A, ATG5, ATG7, ATG12*), it increased autophagic flux through suppression of mTOR phosphorylation at Ser2448. Given that NPNT is a miR-378a target and a key glomerular basement membrane component, we investigated its role in autophagy regulation. NPNT knockdown reduced *ATG2A*, *ATG7*, and *BCN1* expression, but paradoxically increased autophagic flux, independent of mTOR, accompanied by enhanced ERK1/2 phosphorylation. These findings reveal a dual-layered regulatory network in which miR-378a promotes autophagy via mTOR inhibition, whereas NPNT modulates autophagy probably through MAPK-dependent signaling. Our results highlight the complex interplay between miRs, extracellular matrix components, and intracellular signaling pathways in podocyte autophagy. Dysregulation of these pathways in kidney disease may reflect both adaptive and maladaptive responses, providing mechanistic insights and potential therapeutic targets to preserve glomerular filtration barrier integrity in immune-mediated kidney disease.

## Introduction

Autophagy is a major homeostatic and quality control mechanism to maintain cellular integrity being responsible for the bulk degradation of long-lived cytosolic proteins and organelles. Constitutive autophagy has been shown to function as a cell-repair mechanism [1]. Accumulating evidence supports the indispensable role of autophagy in the maintenance of podocyte homeostasis [1-5]. Podocytes exhibit a high basal level of autophagy with abundant autophagosomes [1]. Glomerular injury triggers autophagy to prevent the progression of glomerular disease. Podocyte-specific deletion of class III PI3K vacuolar protein sorting 34, which regulates autophagy, caused proteinuria, glomerulosclerosis, and renal failure [1]. Podocyte specific deletion of *ATG5* resulted in accelerated diabetes-induced podocytopathy with glomerulosclerosis [4]. Dysregulation of autophagy homeostasis is proposed to play a role in the pathophysiology of MGN [3, 6]. In addition to podocytes, autophagy is also important in proximal tubular cells. Proximal tubular cells consume a lot of energy during electrolyte reabsorption, which requires high lysosomal activity and mitochondrial turnover [7]. Thus, impaired autophagy might be one of the common mechanisms underlying podocyte and tubular injury in MGN.

Although, sublytic C5b-9 membrane attack complex was reported to increase autophagic induction, and trigger lysosomal membrane permeabilization, resulting in the accumulation of autophagosomes and autophagic inactivation [3] other potential mechanisms causing autophagy dysfunction in MGN remain poorly understood.

Most publications on the role of autophagy in glomerular disease only assessed autophagy via marker expression levels, such as microtubule-associated protein 1 light chain 3 (LC3). During autophagy, LC3-I is conjugated to phosphatidylethanolamine to form LC3-II, which is localized to isolation membranes and autophagosomes. LC3 levels increase with autophagy induction due to increased autophagosome formation, but also decrease as these forming autophagosomes are turned over. An increase in LC3-II could therefore mean the cell has increased autophagy because there are more autophagosomes, but it could also mean that the cell has decreased autophagy because of inhibited degradation of autophagosomes [8].

In the last years, the importance of microRNAs (miRs) in regulating important signaling pathways and cellular functions became evident [7].

Lately, we investigated the role of miRs as a source of intercompartmental cell-cell communication in the pathogenesis of MGN. We found podocyte-derived miR-378a-3p (miR-378a) is upregulated in patients with MGN and regulates expression and secretion of glomerular extracellular matrix protein nephronectin (NPNT). Overexpression of miR-378a in zebrafish induced edema, proteinuria, loss of podocyte markers and podocyte effacement [9]. In addition to NPNT, glomerular cell derived miR-378a and miR-192 have autophagy associated mRNAs as common targets, according to miR databases and findings in other cell types [9-11]. However, it is unclear how MGN associated miRs regulate podocyte autophagy. Moreover, recent work proposed an active and dynamic signaling role for extracellular matrix-evoked autophagic regulation [12, 13]. Nevertheless, the relationship between the extracellular protein NPNT and autophagy has not been investigated so far.

Here, we aimed to investigate the role of MGN associated miR-378a in the regulation of autophagy in podocytes. We used autophagy flux assay to capture the dynamics of autophagy in cell culture and to be able to differentiate between increase in autophagy and decrease in autophagosome degradation. We further started to unravel the signaling pathways involved in the NPNT-mediated modulation of autophagy.

## Results

### MGN associated miR-378a does not regulate autophagy related genes directly

Given the emerging link between podocyte injury and autophagy dysregulation, we sought to determine whether miR-378a might directly modulate components of the autophagic machinery. *In silico* target prediction analysis identified *ATG2A* and *ATG12* as potential targets of miR-378a (**Supplementary Fig. 1**), both of which encode essential regulators of autophagosome formation and elongation. To experimentally validate these findings, we performed quantitative PCR (qPCR) analyses in cultured human podocytes following transient overexpression of miR-378a. Careful selection of appropriate controls was critical for these experiments. Initially, we included untransfected cells, Lipofectamine-only treated cells, and cells transfected with a non-targeting control miR (miR-Ctrl) in combination with Lipofectamine. However, comparative analysis revealed significant differences in the expression of autophagy-related genes among untransfected cells, Lipofectamine-only treated cells, and miR-Ctrl/Lipofectamine-treated cells (**Supplementary Fig. 2**). These findings are consistent with previous reports demonstrating that control siRNA complexed with Lipofectamine 2000 can itself alter basal autophagy signaling [14].

Based on these observations, we concluded that untransfected cells and Lipofectamine-only treated cells do not provide reliable controls in the context of autophagy-related readouts. Instead, miR-Ctrl transfected with Lipofectamine was selected as the most appropriate comparator for miR-378a/Lipofectamine and miR-192/Lipofectamine transfection experiments, as it controls for both transfection reagent exposure and non-specific miR effects.

Using this optimized experimental setup, we assessed mRNA expression levels of key autophagy-related genes following miR-378a overexpression in human immortalized podocytes. Notably, transfection of podocytes with miR-378a did not result in significant changes in *ATG2A, ATG5, ATG7*, or *ATG12* transcript levels compared with miR-Ctrl (**Fig. 1**). These results suggest that, despite *in silico* prediction of *ATG12* and *ATG2A* as potential targets, miR-378a does not exert a measurable regulatory effect on the transcriptional abundance of these core autophagy genes in cultured human podocytes under the conditions tested.

**Fig. 1.**
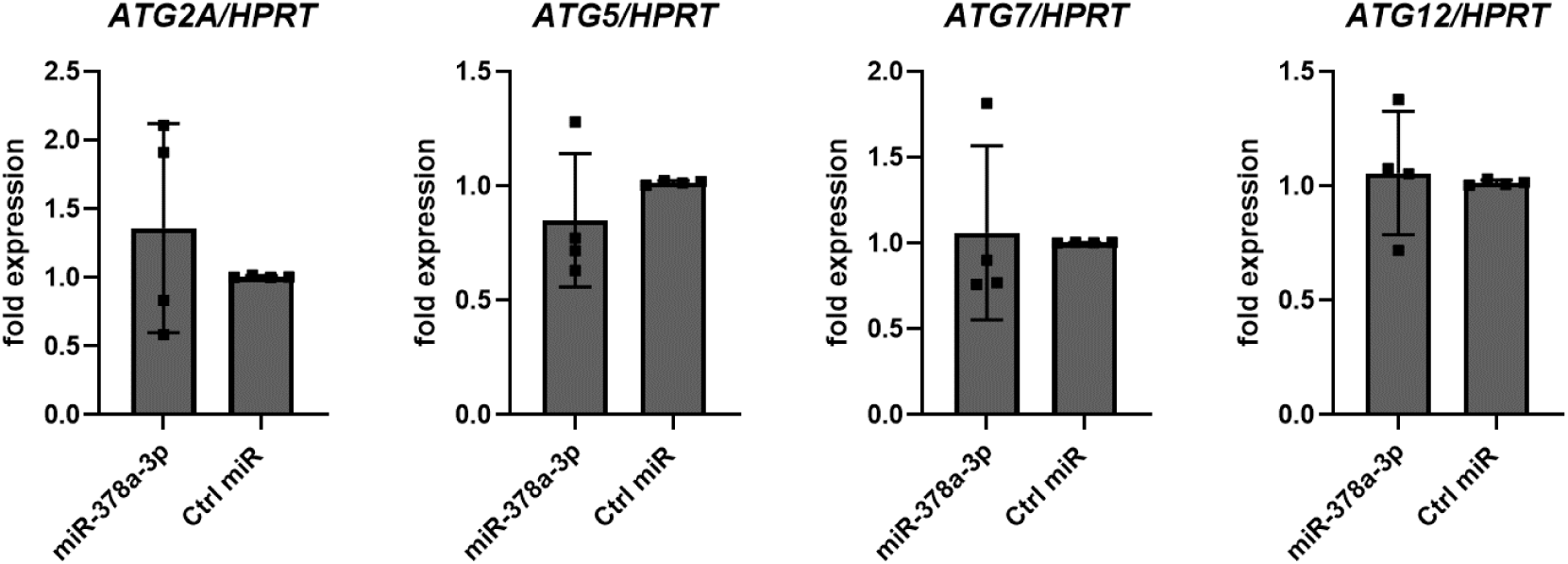
miR-378a does not directly alter transcription of core autophagy genes in podocytes. Quantitative PCR analysis of *ATG2A*, *ATG5*, *ATG7*, and *ATG12* mRNA in human podocytes transfected with miR-378a mimic or miR-Ctrl, 44 h after transfection. mRNA expression was normalized to *HPRT* and given as fold change compared to control transfected cells. n = 4 independent experiments.

### miR-378a enhances autophagic flux in podocytes

Interpretation of static LC3 measurements is inherently limited, as elevated LC3 levels may reflect either increased autophagosome biogenesis or impaired autophagosome clearance due to defective lysosomal fusion, resulting in accumulation of undegraded LC3-positive vesicles. Consequently, assessment of single, steady-state autophagy markers is insufficient to accurately quantify autophagic activity, which represents a dynamic and highly regulated process. To rigorously evaluate the impact of miR-378a on autophagy, we therefore focused on measuring autophagic flux rather than relying on static endpoints.

Autophagic flux encompasses the complete autophagy pathway, including autophagosome initiation, elongation, maturation, fusion with lysosomes, lysosomal degradation of cargo, and recycling of breakdown products into the cytosol. Experimentally, flux can be quantified by comparing LC3-II expression in the presence and absence of pharmacologically blocked autophagosome-lysosome fusion, thereby enabling accumulation of newly formed autophagosomes and allowing discrimination between enhanced formation and impaired degradation.

To this end, we utilized bafilomycin A1 (BafA), an inhibitor of vacuolar H⁺-ATPase, which prevents lysosomal acidification and autophagosome–lysosome fusion. Prior to performing functional experiments, we conducted dose-optimization studies to determine a concentration that reliably inhibited autophagosome-lysosome fusion in cultured human podocytes without inducing nonspecific toxicity. Based on these analyses, 30 nM BafA was selected as the optimal concentration for subsequent flux measurements (**Supplementary Fig. 3**). Using this validated assay, we next investigated the effect of miR-378a on autophagic flux. Overexpression of miR-378a via transfection with a synthetic mimic significantly increased autophagic flux in human podocytes, as evidenced by enhanced accumulation of LC3-II in the presence of BafA, compared to miR-Ctrl-transfected cells (**Fig. 2a**). A comparable increase in autophagic flux was observed in HK-2 tubular epithelial cells following miR-378a overexpression (**Supplementary Fig. 4a**), indicating that miR-378a-mediated autophagy regulation is conserved across renal cell types. In contrast, inhibition of endogenous miR-378a using a specific antisense inhibitor significantly reduced autophagic flux in cultured human podocytes (**Fig. 2b**).

**Fig. 2.**
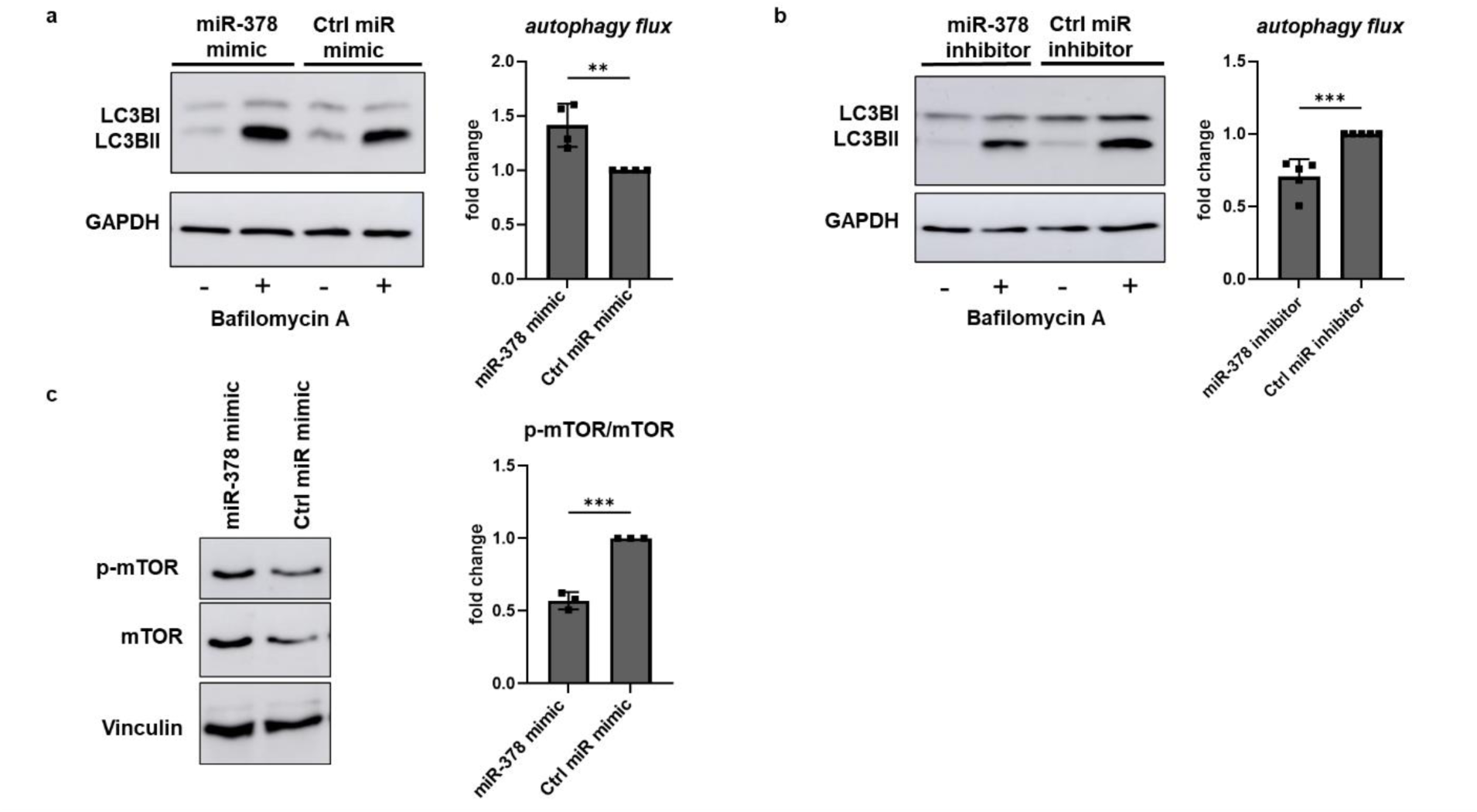
miR-378a enhances autophagic flux via mTOR inhibition in podocytes. a. Autophagic flux assay by western blot analysis. Cultured human podocytes were transfected with miR-378a mimic or miR-Ctrl and treated with 30 nM bafilomycin A1 (BafA) for 46 h to block autophagosome-lysosome fusion or left untreated. LC3-II protein accumulation was assessed by Western blot. Autophagic flux was calculated by subtracting basal LC3-II levels from LC3-II levels after BafA treatment within each group. Protein expression was normalized to GAPDH. Autophagy flux is given as fold change compared to miR-Ctr, n = 4 independent experiments b. Loss-of-function analysis. Inhibition of endogenous miR-378a using a specific antisense inhibitor decreased autophagic flux in podocytes, confirming a positive regulatory role for miR-378a in autophagy. n =4 independent experiments, c. Western blot analysis of mTOR phosphorylation at Ser2448 in podocytes following miR-378a mimic or miR-Ctrl transfection. Phospho-mTOR/total mTOR ratio was normalized to Vinculin and given as fold change compared to miR-Ctrl. n = 3 independent experiments, ** *p* < 0.01; *** *p* < 0.001.

Taken together, these complementary gain- and loss-of-function experiments demonstrate that miR-378a positively regulates autophagic flux in podocytes. Importantly, the use of a flux-based approach confirms that miR-378a enhances autophagosome formation and turnover rather than merely causing accumulation of undegraded autophagic vesicles, thereby establishing miR-378a as a functional modulator of autophagy in renal epithelial cells.

### Upregulated autophagy flux by miR-378a is mediated by mTOR pathway

After the observation that miR-378a enhances autophagic flux in podocytes, we next sought to delineate the upstream signaling pathway responsible for this effect. The mammalian target of rapamycin (mTOR) is a central negative regulator of autophagy and integrates nutrient, energy, and growth factor signals to suppress autophagosome initiation under nutrient-replete conditions [15]. Inhibition of mTOR complex 1 (mTORC1) activity is a well-characterized trigger for autophagy induction, primarily through de-repression of the ULK1 complex and subsequent activation of the autophagic machinery.

To determine whether miR-378a modulates autophagy through mTOR signaling, we assessed mTOR activity by quantifying phosphorylation at Ser2448, a commonly used marker of mTORC1 activation. Western blot analysis revealed a significant reduction in the phospho-mTOR (Ser2448)/total mTOR ratio following transfection of podocytes with a miR-378a mimic compared with miR-Ctrl-transfected cells (**Fig. 2c**). This decrease in mTOR phosphorylation indicates suppression of mTORC1 activity and is consistent with enhanced autophagy initiation. These findings support a model in which miR-378a promotes autophagic flux by functionally inhibiting mTOR signaling, thereby relieving its tonic suppression of autophagosome formation in podocytes. Importantly, similar reductions in mTOR phosphorylation were observed in HK-2 tubular epithelial cells following miR-378a overexpression (**Supplementary Fig. 4b**), suggesting that miR-378a-mediated modulation of mTOR activity represents a conserved regulatory mechanism across renal cell types.

Collectively, these data identify mTOR as a critical signaling node downstream of miR-378a and provide mechanistic insight into how miR-378a enhances autophagic flux in podocytes.

### NPNT knockdown modulates autophagy gene expression

Building on our previous work demonstrating the regulatory role of podocyte miR-378a on NPNT and its impact on glomerular homeostasis [9], we next sought to determine whether NPNT itself influences autophagy-related gene expression and functional autophagic activity. NPNT is an essential extracellular matrix protein localized in the glomerular basement membrane, and emerging evidence suggests that perturbations in GBM composition can affect podocyte stress responses and intracellular signaling pathways. To investigate the direct contribution of NPNT to autophagy regulation, we transfected cultured human podocytes with NPNT-specific siRNA or non-targeting control siRNA and assessed mRNA expression levels of key autophagy-related genes. Efficient knockdown of *NPNT* was confirmed at the transcript level (**Fig. 3a**). Notably, *NPNT* knockdown resulted in a significant downregulation of *ATG2A*, *ATG7*, and *BCN1* transcripts (**Fig. 3b**), suggesting that NPNT positively modulates the expression of core autophagy machinery components.

**Fig. 3.**
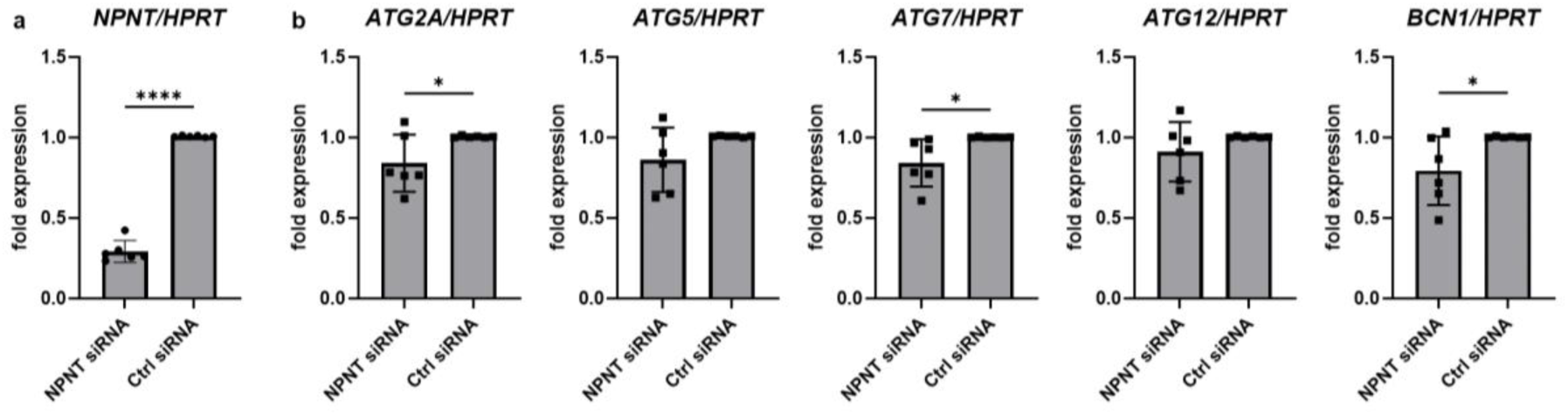
NPNT knockdown decreases autophagy-related gene expression in podocytes. Quantitative PCR analysis of *NPNT* (a) as well as *ATG2A*, *ATG5, ATG7*, *ATG12* and *BCN1* (b) *mRNA* expression of cultured human podocytes transfected with NPNT-specific siRNA (NPNT siRNA) or non-targeting control siRNA (Ctrl siRNA). Data as normalized to *HPRT* and given as fold change compared to Ctrl siRNA. n = 6 independent experiments, * *p* < 0.05; **** *p* < 0.0001.

### NPNT knockdown increases autophagy flux

Surprisingly, despite the observed reduction in autophagy-related mRNAs, functional assessment of autophagic flux as described above, revealed an increase in autophagy following NPNT knockdown. Using western blot analysis to quantify LC3-II accumulation in the presence and absence of lysosomal inhibition, we observed enhanced autophagic flux in NPNT-depleted podocytes compared with control cells (**Fig. 4a**). This paradoxical effect indicates that, although NPNT supports basal expression of ATG genes, its loss triggers compensatory mechanisms which stimulate autophagosome formation and turnover, potentially as a cellular stress response to altered GBM interactions. Interestingly, when we stimulated cultured podocytes with NPNT protein, we detected the opposite effect, with decreased autophagy flux (**Fig. 4b**), which underlines the significance of NPNT levels for flux modulation.

**Fig. 4.**
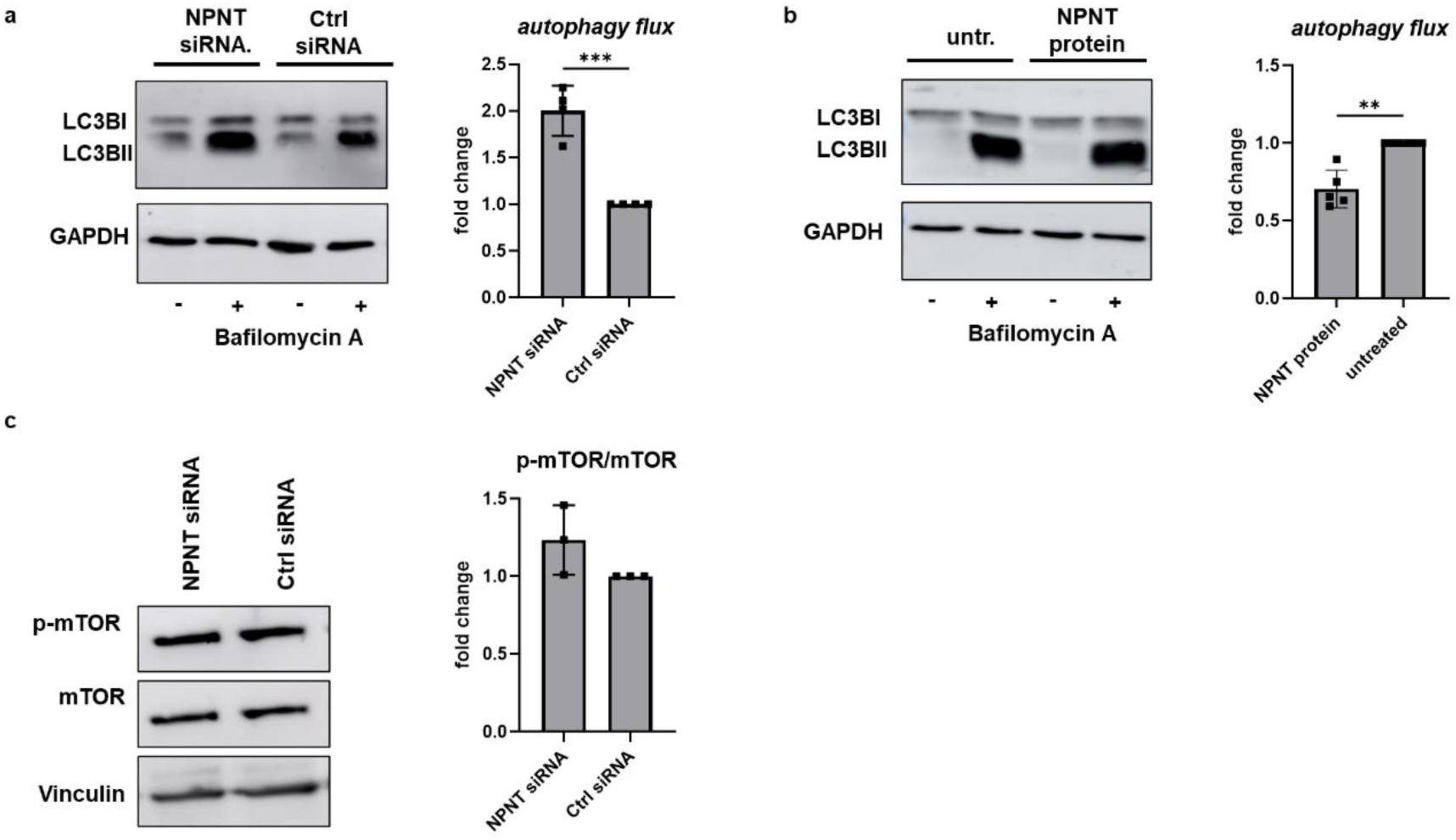
NPNT depletion increases autophagic flux independent of mTOR in podocytes. a. Autophagic flux analysis of cultured human podocytes after transfection with NPNT-specific siRNA (NPNT siRNA) or non-targeting control siRNA (Ctrl siRNA) +/- 30 nM BafA for 44 h to block autophagosome-lysosome fusion. LC3-II protein accumulation was assessed by Western blot. Autophagic flux was calculated by subtracting basal LC3-II levels from LC3-II levels after BafA treatment within each group. Protein expression was normalized to GAPDH. Autophagy flux is given as fold change compared to Ctr siRNA, n = 4 independent experiments. b. Cells as in a. were stimulated with NPNT protein or unstimulated +/- 30 µM BafA Autophagic flux was assessed as in a., n = 5 independent experiments. c. Western blot analysis of mTOR phosphorylation at Ser2448 in podocytes following NPNT siRNA or Ctrl siRNA transfection. Phospho-mTOR/total mTOR ratio was normalized to Vinculin and given as fold change compared to miR-Ctrl. n = 3 independent experiments. ** *p* < 0.01 *** *p* < 0.01

As we identified mTOR pathway as a mediator of miR-378a induced autophagy flux, we also investigated this pathway after siRNA induced NPNT knockdown. However, phosphorylated mTOR to mTOR levels were unchanged (**Fig. 4c**).

NPNT consists of three main domains, the N-terminal EGF-like repeats, the linker-region containing two integrin-binding motifs, and the C-terminal MAM domain [16]. The EGF-like repeats may bind the epidermal growth factor receptor (EGFR). Pro- as well as anti-autophagy effects of EGFR signaling were reported before [17, 18]. To investigate a possible mechanism for NPNT-mediated modulation of autophagy, we focused on components of the EGRF pathway. Phosphorylation of possible pathway candidates was analyzed in immortalized podocytes after transfection with NPNT siRNA or Ctrl siRNA. No significant difference was observed in pEGFR/EGFR (**Fig. 5a**) and pAKT/AKT (**Fig. 5b**) ratios. However, the ratio of pERK1/2 was increased after NPNT siRNA treatment (**Fig. 5c**), hinting that the effects on autophagy are not directly mediated by EGFR.

**Fig. 5.**
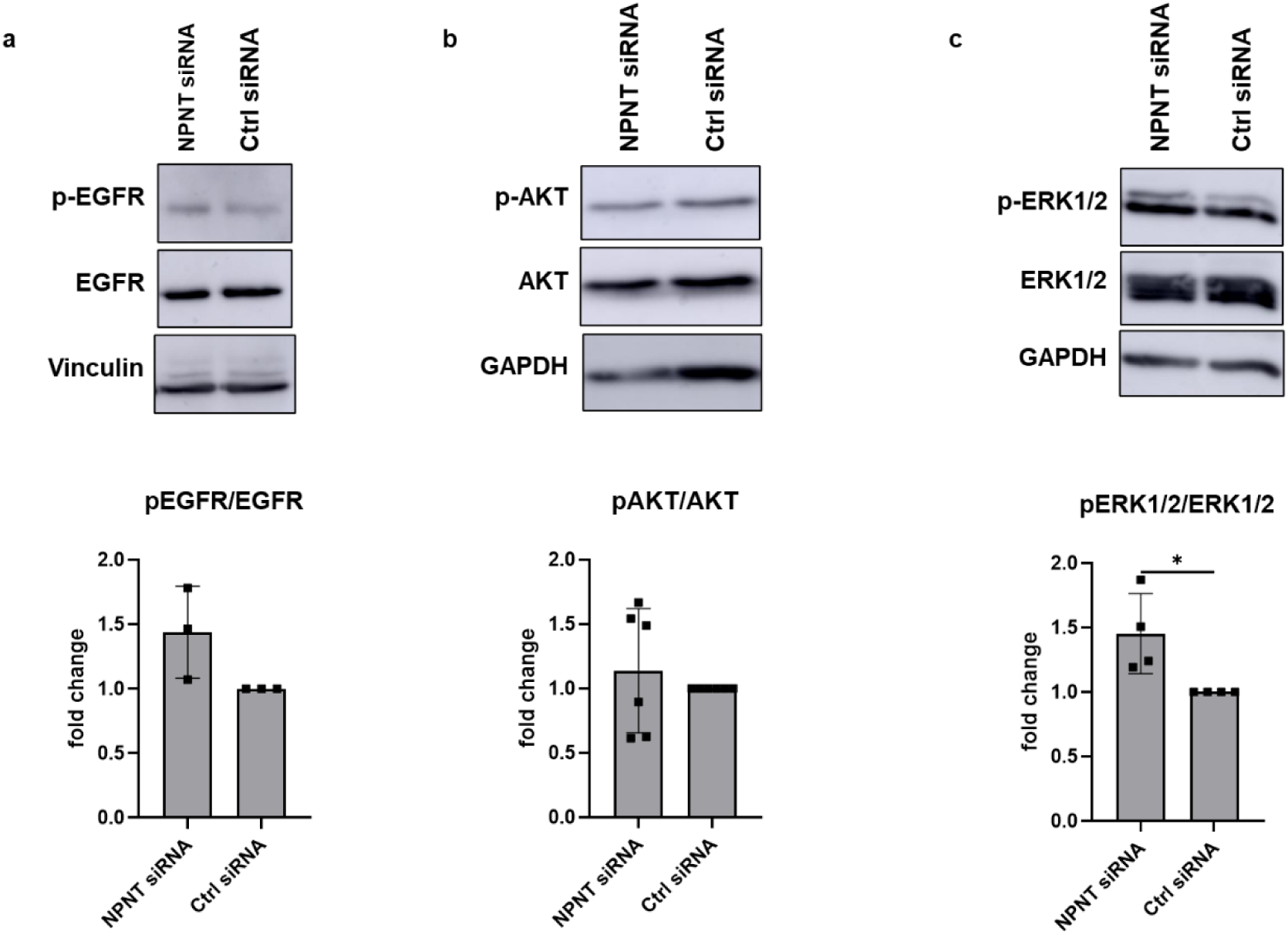
siRNA-mediated NPNT depletion increases ERK signaling in podocytes. a-c: Western blot analysis of EGFR (a), AKT (b) and ERK1/2 (c) phosphorylation in podocytes following NPNT siRNA or control siRNA transfection. Phospho-protein levels were normalized to total protein levels, respectively. Vinculin (a) or GAPDH (b, c) served as loading controls. Data is presented as fold change relative to control siRNA (Ctrl). Data represent mean ± SD, n = 3-5 independent experiments. * p <0.05.

These findings reveal a complex link between NPNT expression and podocyte autophagy, where NPNT regulates transcription of autophagy-associated genes, yet its temporal depletion paradoxically promotes autophagic flux. This dual effect underscores the intricate crosstalk between extracellular matrix composition and intracellular proteostatic mechanisms, providing a mechanistic link between GBM perturbation and adaptive autophagy responses in podocytes. As NPNT has been described before to bind to the RGD motif of integrins, this family of transmembrane proteins imposes itself as plausible components to mediate extracellular NPNT signals to modulate autophagy in podocytes.

## Discussion

An autophagy-dependent dual role with protective or detrimental effects has been identified in ischemia-reperfusion renal injury and protein overload-associated tubular epithelial cell lesions [19] highlighting the importance of tight regulation of autophagy. *In vitro* experiments suggested that autophagic pathways play key roles for the activation of this self-protection mechanism of the cell. Podocytes and tubular cells are dependent on a relatively high level of autophagy [7]. Impaired autophagy may be involved in the pathogenesis of podocyte loss, leading to massive proteinuria in diabetic nephropathy [4, 20].

Autophagy is also disturbed in MGN [3, 6]. A pronounced upregulation of LC3 in podocytes adjacent to the GBM in MGN patient biopsies was observed, confirming dysregulated autophagy signaling in human disease and establishing a spatial link between immune-mediated GBM injury and activation of the autophagic machinery. Importantly, the punctate LC3 pattern observed is consistent with increased autophagosome formation, although static LC3 measurements cannot discriminate between enhanced autophagy initiation and impaired autophagosome turnover [3]. *Jin et al.* found that autophagy contributed to podocyte injury in MGN and the number of autophagosomes in podocytes was related to the pathological classification [21]. Furthermore, an upregulation of lysosomal proteins, especially of Limp2 [22] and cathepsin D [23] is prominent in MGN. However, the underlying mechanism of impaired autophagy in MGN needs further investigations. Autophagy might play a complex, dual role in MGN. While it protects podocytes from injury and apoptosis, it may also facilitate the presentation of podocyte antigens, contributing to autoantibody production and immune complex formation [24].

Our study provides new mechanistic insights into the regulation of podocyte autophagy and highlights the coordinated interplay between miRs, extracellular matrix proteins, and intracellular signaling pathways. Podocytes are uniquely dependent on efficient autophagic flux due to their highly specialized, terminally differentiated state and limited regenerative capacity. Both podocytes and tubular epithelial cells rely on relatively high basal levels of autophagy to maintain cellular homeostasis, and impaired autophagy has been implicated as a common mechanism underlying glomerular and tubular injury [7, 19].

To investigate molecular drivers of this dysregulation, we focused on miR-378a, which we previously found to be upregulated in podocytes and urine from patients with MGN [9]. *In silico* analyses suggested potential targeting of autophagy genes such as *ATG12* and *ATG2A*; however, overexpression or inhibition of miR-378a in cultured podocytes did not significantly alter ATG transcript levels, indicating that miR-378a does not directly regulate the transcription of core autophagy machinery. Notably, static measurements of LC3 alone are insufficient to capture the dynamic nature of autophagy, as LC3-II accumulation may reflect either increased autophagosome formation or impaired degradation [21-23]. Therefore, we assessed autophagic flux, a more accurate measure that integrates autophagosome formation, lysosomal fusion, and turnover. Using bafilomycin A1 to block autophagosome-lysosome fusion, we found that miR-378a overexpression robustly increased autophagic flux in podocytes and HK-2 tubular epithelial cells, whereas inhibition of miR-378a reduced flux. These results demonstrate that miR-378a positively regulates autophagy functionally, independent of direct transcriptional effects on ATG genes. Mechanistically, miR-378a suppressed mTOR phosphorylation at Ser2448, a central negative regulator of autophagy [15, 25], thereby relieving mTOR-mediated inhibition of autophagosome formation. The conservation of this effect in tubular cells underscores the broader relevance of miR-378a in renal epithelial autophagy regulation.

MTOR was shown to regulate autophagic flux in the glomerulus [26], but we are the first to identify MGN associated miR-378a as an upstream post-transcriptional regulator of mTOR phosphorylation in human podocytes. Interestingly, podocytes maintain high basal levels of autophagy independent of mTOR signaling [5].

Given the role of NPNT as a miR-378a target and its localization to the GBM [9], we examined whether NPNT itself influences autophagy We could show that NPNT knockdown enhances autophagic flux despite reducing the mRNA expression of several canonical autophagy-related genes (*ATG2A, ATG7*, and *BCN1*) and without suppressing mTOR activity.

While EGFR represents a prototypical receptor tyrosine kinase activating ERK in response to soluble growth factors, we did not observe alterations in EGFR phosphorylation, nor in downstream AKT activation. Instead, the selective increase in ERK1/2 phosphorylation points toward an alternative mTOR- and probably EGFR-independent regulatory axis controlling autophagic dynamics in podocytes. Given that NPNT is an extracellular matrix protein and ligand for integrins, its depletion is expected to perturb cell–matrix interactions rather than growth factor receptor engagement. While NPNT contains EGF-like repeats capable of binding EGFR [16], it has also previously been described to bind to integrins via its RGD-binding motif [27]. Integrins are a superfamily of cell adhesion receptors, which bind to soluble ligands and ligands both in the ECM and on the cell surface. Upon binding of an extracellular ligand, integrins transduce the signal intracellularly, which trigger changes in cell behavior, such as adhesion, survival, gene expression or motility [28]. Integrins, autophagy and altered ERK1/2 activation have recently been described to be involved in detachment-induced autophagy, a process called anoikis [29]. Thus, the observed ERK activation likely reflects matrix-sensing signaling rather than growth factor-driven stimulation. These findings support a model in which autophagy regulation in podocytes is governed by multilayered signaling networks which integrate extracellular matrix composition, MAP kinase signaling, and miR-mediated post-transcriptional control. Together, our data expand the current understanding of autophagy regulation in the glomerulus by identifying a dual regulatory system: miR-378a-mediated suppression of mTOR activity and NPNT-dependent modulation of autophagy flux, possibly mediated by integrin signaling (**Fig. 6**). This layered control may be particularly relevant in immune-mediated kidney diseases, where extracellular matrix alterations and inflammatory signaling converge on podocyte stress responses.

**Fig. 6.**
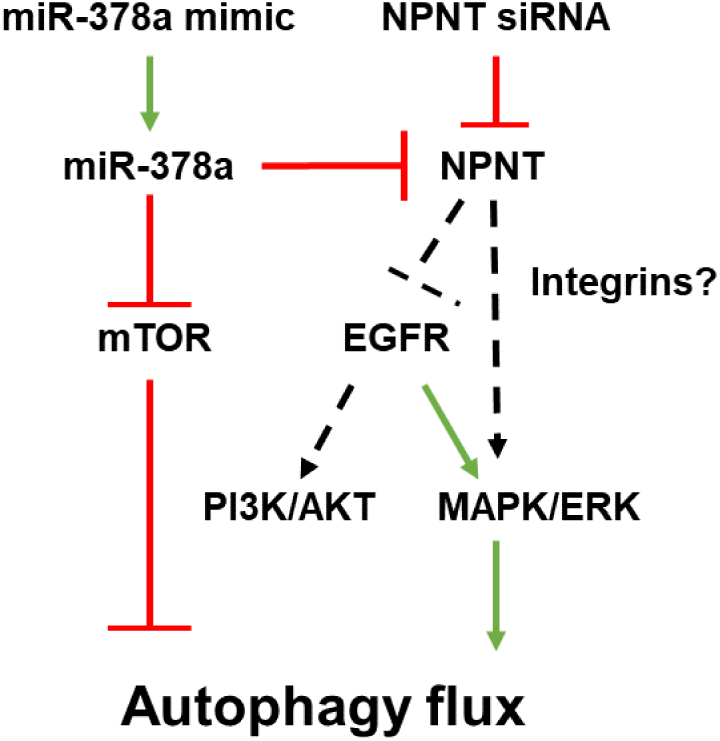
Different pathways for altering autophagy flux involving NPNT modulation. Schematic depiction of pathway hypothesis for alteration of autophagy flux following alterations in NPNT expression.

This dual regulation underscores the intricate crosstalk between extracellular matrix composition, miR signaling, and intracellular proteostatic mechanisms, providing a mechanistic framework linking GBM perturbation, miR dysregulation, and adaptive autophagy responses. The dynamic, context-dependent nature of autophagy emphasizes the need for precise regulation, as both insufficient and excessive autophagic activity may contribute to podocyte injury and progression of glomerular disease [19, 21].

## Conclusion

In conclusion, our work establishes miR-378a as a functional enhancer of podocyte autophagy, identifies NPNT as a modulatory extracellular matrix component, and uncovers mechanistic links to mTOR and MAPK signaling. These findings not only advance our understanding of autophagy dysregulation in MGN, but also highlight potential therapeutic targets to restore proteostatic balance and protect the glomerular filtration barrier in immune-mediated kidney disease. Thus, miRNAs represent an additional layer in the intricate interconnection between autophagy and extracellular matrix. Comprehensive knowledge of miRs and related networks modulating autophagy might contribute to innovative treatment approach of renal diseases in the future.

## Methods

### Cell culture

Conditionally immortalized human podocytes were used, which were initially created by Saleem et al. [30] and now represent a well-established cell culture model in podocyte research around the world. This podocyte cell line proliferates under permissive conditions at 33 °C. When cultivated at 37 °C, the SV40 T-antigen is inactivated for cell differentiation (8 to 10 days). The culture medium for human podocytes was RPMI 1640 Medium (Gibco, via Thermo Fisher Scientific, Waltham, MA, USA) with 10% fetal calf serum (FCS; PAN-Biotech, Aidenbach, Germany), 1% Penicillin/Streptomycin, and 0.1% Insulin-Transferrin-Selenium (Gibco).

As proximal tubular cell line HK-2 cells were used, which are derived from an adult male human kidney. HK2 cells were cultured in DMEM/F12 1:1 media (Gibco), supplemented with 5% FCS (PAN-Biotech) and 1% Penicillin/Streptomycin, at 37° C, 5% CO_2_.

### miRNA/siRNA transfection

Podocytes were differentiated for 8 to 10 days and were transfected with 5µM of mirVana® hsa-miR-378-3p mimic (assay ID: MC11360), mirVana® miRNA inhibitor (assay ID: MH11360), mirVana^TM^ miR inhibitor Negative Control #1, mirVana™ miRNA Mimic, Negative Control #1 (4464059), Silencer™ Select Negative Control #1, Silencer™ Select NPNT siRNA, (assay ID 128582, all Thermo Fisher Scientific). Transfection was carried out as published before [9]. with Lipofectamine 2000 in Opti-MEM^TM^ (both Thermo Fisher Scientific).

### Bafilomycin treatment

Bafilomycin A1 (BafA, Sigma-Aldrich) was used for autophagy flux assays. BafA was added to cells at a concentration of 30 nM for podocytes and 50 nM for HK-2 cells. In combination with transfection, BafA was added with the media after 24 hours of transfection. Stimulation times varied between cell types and analyzed targets. Specific incubation times are given in the figure legends.

### qPCR

Whole cell lysate RNA was isolated by using the ReliaPrep™ RNA Miniprep System (Promega, Madison, WI, USA), according to the protocol provided by the manufacturer. Reverse transcription of RNA was done by using the following reagents: M-MLV RT 50.000 U in 5x RT buffer (Promega), dNTP Mix (Promega), random hexamer primer (ThermoFisher Scientific), and RiboLock (Thermo Fisher Scientific). Reverse transcription protocol: 25 °C for 5 min, 40 °C for 60 min, and 70 °C for 10 min. For quantitative real-time PCR SYBR Green dye (Maxima SYBR Green/ROX qPCR Master Mix, Thermo Fisher Scientific) was used and run with the following protocol: 10 min at 95 °C followed by 40 cycles of 15 s at 95 °C and 1 min at 60 °C, followed by 15 s at 95 °C, 1 min at 60 °C and 15 s at 95 °C. Each sample was run in triplicates. The following primers were used, table 1:

**Table 1:**
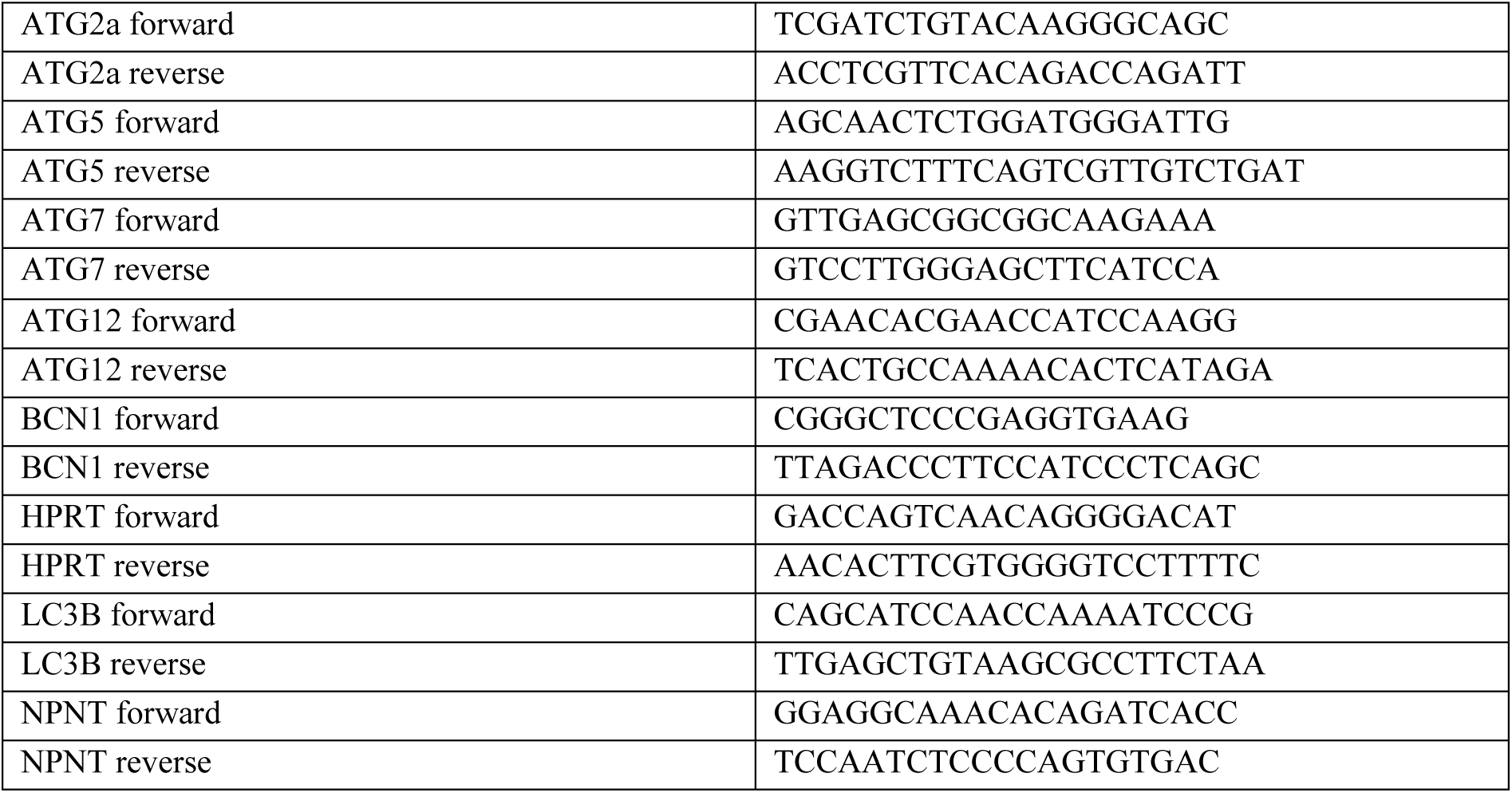
Primer Sequence. Analysis of qPCR results was done by calculating ΔΔCt values and normalization to *HPRT*.

### Western blot

Cells were harvested by lysing in RIPA buffer containing protease and phosphatase inhibitors. 20 μg of podocyte cell lysate were loaded on 8 %, 10 % or 15 % SDS-PAGE, depending on target size, and transferred to nitrocellulose or PVDF (phospho-antibody blots) membranes by wet blot transfer. Detection of protein bands was performed using horseradish peroxidase-labelled secondary antibodies and visualized using enhanced chemiluminescence reagents. Primary antibodies used in this study:

Phospho-Akt (Ser473, #9271S, Cell Signaling), AKT (#9272, Cell Singaling), phospho-EGFR (Tyr1068, #3777S, Cell Signaling), EGFR (#2232, Cell Signaling), phospho-ERK1/2 (phosphor-p44/42 MAPK (Thr202/Tyr204), #9106, Cell Signaling), ERK1/2 (p44/42 MAPK, #9102, Cell Signaling), GAPDH (sc-32233, Santa Cruz), LC3 (NB100-2220, Novus), phospho-mTOR (Ser2448, #5536S, Cell Signaling), mTOR (2983S, Cell Signaling), vinculin (V9131, Sigma-Aldrich). Secondary antibodies: Polyclonal Goat anti-mouse Immunoglobulins/HRP, Polyclonal Goat anti-rabbit Immunoglobulins/HRP (Agilent Technologies Denmark)).

Phospho-antibody-secondary antibody complexes were stripped off the membrane using ReBlot Plus Mild Antibody Stripping Solution, 10x (Sigma-Aldrich) according to the manufacturers’ protocol. Successful stripping was tested by incubation with secondary antibody only and subsequent development of chemiluminescent signal. Absence of signal was considered successful stripping.

Semi-quantitative analysis of protein expression was performed using ImageJ software (National Institutes of Health, Bethesda, MD) by measuring band intensities of target proteins and a reference protein as indicated. Fold expression was calculated. For autophagy flux calculation, signal ratio of LC3 II/GAPDH after incubation with BafA minus LC3 II/GAPDH ratio without BafA was determined. For pooling of independent experiments, the fold change of autophagy flux was calculated.

### Statistical analysis

Data is shown as mean ± SD and were compared by ANOVA or students t-test to test for statistical significance. Statistical analysis was performed with GraphPad Prism software.

## Abbreviations

AKT: protein kinase b
ATG: autophagy-related gene
ATG2a: autophagy-related gene 2a
ATG 5: autophagy-related gene 5
ATG 7: autophagy-related gene 7
ATG 12: autophagy-related gene 12
BAFA/BAFA1: bafilomycin a1
BCN1: beclin 1
c5b-9: complement component 5b-9 (membrane attack complex)
ctrl: control
ECM: extracellular matrix
EGF: epidermal growth factor
EGFR: epidermal growth factor receptor
ERK1/2: extracellular signal-regulated kinases 1 and 2
GBM: glomerular basement membrane
HK-2: human kidney-2 cells
LC3: microtubule-associated protein 1 light chain 3
LC3-i: microtubule-associated protein 1 light chain 3-i
LC3-ii: microtubule-associated protein 1 light chain 3-ii
LIMP2: lysosomal integral membrane protein 2
MAPK: mitogen-activated protein kinase
miR: micro RNA
mir-378a: microRNA-378a
miR-192: microRNA-192
mRNA: messenger RNA
MGN: membranous glomerulonephritis
mTOR: mechanistic target of rapamycin
mTORC1: mechanistic target of rapamycin complex 1
NPNT: nephronectin
qPCR: quantitative polymerase chain reaction
PI3K: phosphoinositide 3-kinase
RGD: arginine–glycine–aspartate motif
siRNA: small interfering RNA
ULK1: unc-51 like autophagy activating kinase 1

## Data availability statement

All datasets are available on request.

## Authors contribution

Conceptualization, N.S., J.M.-D. and N.S.; methodology, N.S., M.W., M.H., A.O.; validation, N.S., A.O. and J.M.-D.; formal analysis, N.S. and J.M.D.; investigation, N.S. and J.M.-D.; resources, M.S. and J.M.-D.; writing—original draft preparation, N.S. and J.M.-D.; writing—review and editing, N.S. and J.M.-D.; visualization, N.S., J-M.-D; supervision, J.M.-D. and N.S.; project administration, J.M.-D.; funding acquisition, J.M.-D. and N.S. All authors have read and agreed to the published version of the manuscript.

