## Supplementary Figure for "miR-378a and NPNT coordinate autophagy regulation in podocytes through mTOR and MAPK signaling"

**Supplements**

**Supplementary Figures**

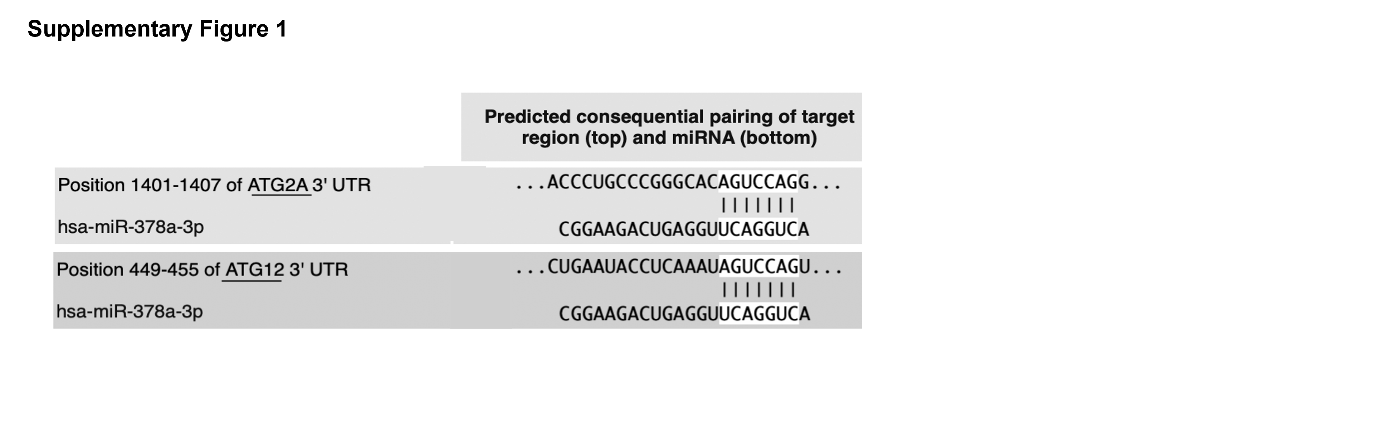

**Supplementary Fig. 1 *miR binding to autophagy mRNAs***

Position and binding site of miR-378a-3p to 3’UTR region of *ATG2A* and *ATG12*.

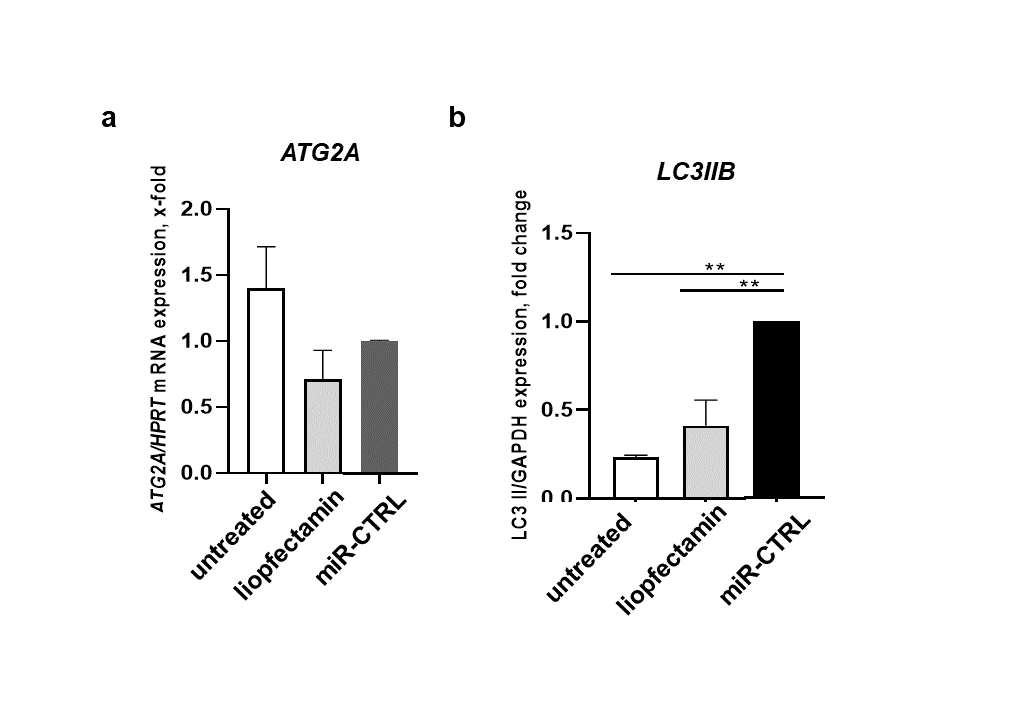

**Supplementary Fig 2. *Effect of Lipofectamine on autophagy***

Quantitative PCR analysis of *ATG2A* (a) and *LC3IIB* (b) mRNA in human podocytes untreated or transfeceted with lipofectamine only (lipofectamine) or (lipofectamine + microRNA CTRL (miR). mRNA expression was normalized to *HPRT* (a) or *GAPDH* (b) and given as fold change normalized to miR-CTRL (*ATG2A*) or untreated (*LC3IIB*). n = 3 independent experiments. ** p<0.01.

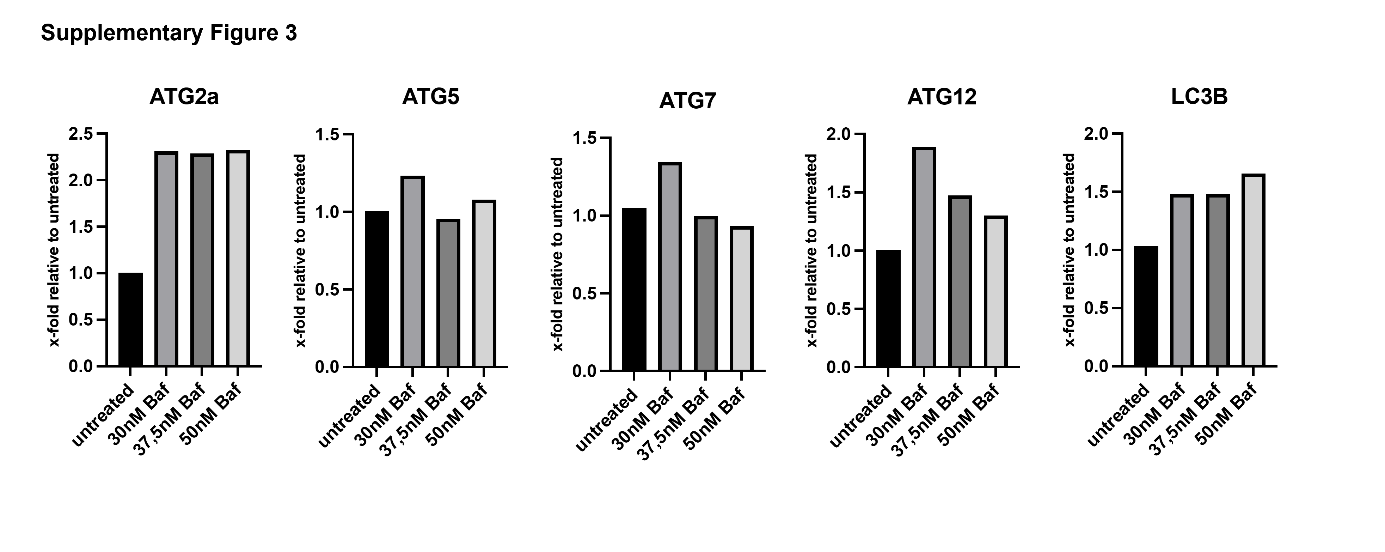

**Supplementary Fig. 3 *Dose finding of Bafilomycin A in cultured human podocytes***

Quantitative PCR analysis of *ATG2A*, *ATG5*, *ATG7*, *ATG12* and *LC3* mRNA in human podocytes treated with different concentrations of Bafilomycin (BafA) or left untreated. mRNA expression was normalized to *HPRT* and given as fold change compared to control transfected cells. n = 1.

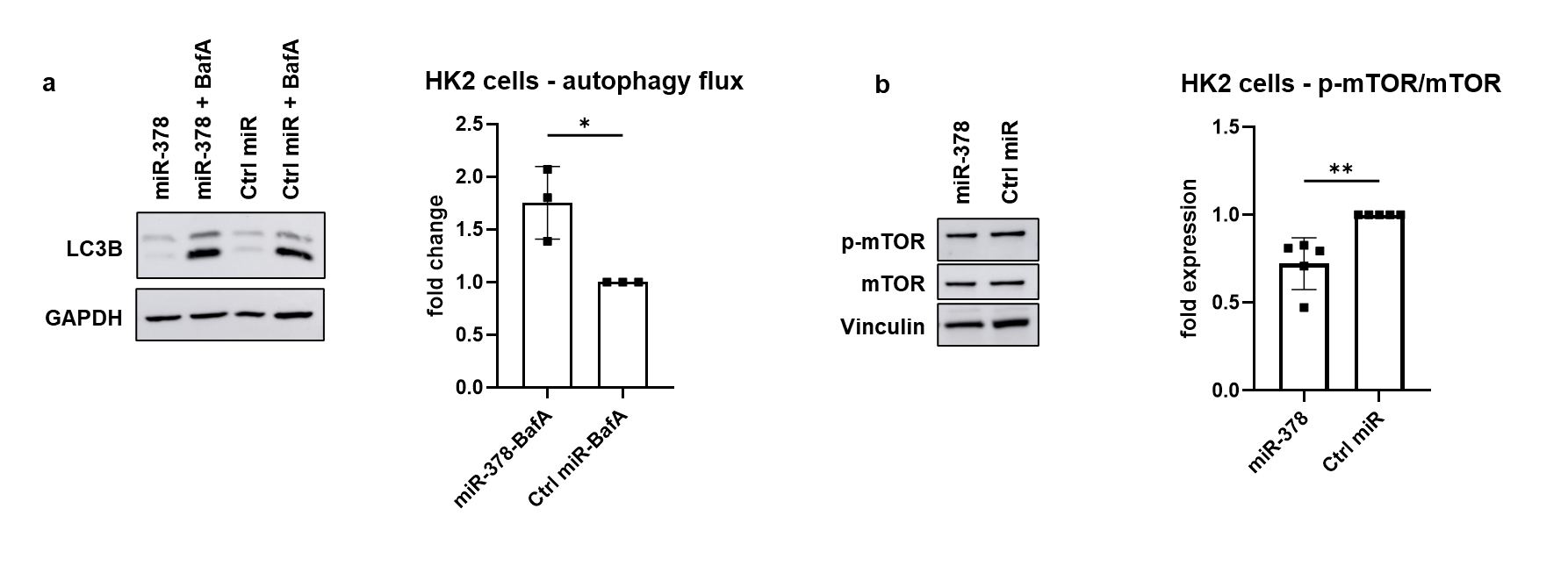

**Supplementary Fig. 4 *miR-378a enhances autophagic flux via mTOR inhibition in HK‑2 cells***

a: Cultured HK-2 cells were transfected with miR-378a mimic or miR-Ctrl and treated with 50 nM BafA for 52 h to block autophagosome-lysosome fusion or left untreated. LC3-II protein accumulation was assessed by Western blot. Autophagic flux was calculated by subtracting basal LC3-II levels from LC3-II levels after bafilomycin A1 treatment within each group. Protein expression was normalized to GAPDH. Autophagy flux is given as fold change compared to miR-Ctr, n = 3 independent experiments, * p < 0.05,

b: Western blot analysis of mTOR phosphorylation at Ser2448 in HK-2 cells following miR-378a mimic or miR-Ctrl transfection. Phospho-mTOR/total mTOR ratio is given as fold change compared to miR-Ctrl. Vinculin was used as loading control. n = 5 independent experiments, ** p < 0.01.
